# Discriminating changes in protein structure using PTAD conjugation to tyrosine

**DOI:** 10.1101/2020.02.04.934406

**Authors:** Mahta Moinpour, Natalie K. Barker, Lindsay E. Guzman, John C. Jewett, Paul R. Langlais, Jacob C. Schwartz

**Affiliations:** Department of Chemistry and Biochemistry, University of Arizona, Tucson, AZ 85721, USA; Department of Medicine, Division of Endocrinology, University of Arizona College of Medicine, Tucson, AZ 85721, USA

**Keywords:** tyrosine, triazolinediones, protein folding, low complexity, conjugation

## Abstract

Chemical modification of proteins has been crucial in engineering protein-based therapies, targeted biopharmaceutics, molecular probes, and biomaterials. Here, we explore the use of a conjugation-based approach to sense alternative conformational states in proteins. Tyrosine has both hydrophobic and hydrophilic qualities, thus allowing it to be positioned at protein surfaces, or binding interfaces, or to be buried within a protein. Tyrosine can be conjugated with 4-phenyl-3H-1,2,4-triazole-3,5(4H)-dione (PTAD). We hypothesized that individual protein conformations could be distinguished by labeling tyrosine residues in the protein with PTAD. We conjugated tyrosine residues in a well-folded protein, bovine serum albumin (BSA), and quantified labeled tyrosine with LC-MS/MS. We applied this approach to alternative conformations of BSA produced in the presence of urea. The amount of PTAD labeling was found to relate to the depth of each tyrosine relative to the protein surface. This study demonstrates a new use of tyrosine conjugation using PTAD as an analytic tool able to distinguish the conformational states of a protein.

## INTRODUCTION

Protein structure is driven by the interactions of the 20 amino acids with solvent and other amino acids^1, 2^. Identifying residues near the protein surface can be an effective approach to distinguish alternative protein structures and conformational states^3–5^. However, the conformational states of many important proteins are a challenge to analyze by standard biophysical techniques, such as those based on X-ray diffraction and nuclear magnetic resonance. The level of detail provided by these techniques is most limited in cases of intrinsically disordered and low sequence complexity proteins^6–8^. Covalent protein modification is a well-established tool used in numerous applications including biomedical research, drug development, and biomaterial engineering^9^. These modifications can also be sensitive to protein structure^10^. The use of chemical modifications in alternative protein conformations offers attractive benefits, such as the preservation of a chemical fingerprint through covalent bonds, which can reveal structural information in analytic methods that require the protein denaturation or digestion by proteases^3, 10, 11^.

Chemical approaches to protein modification have recently been expanded to provide new tools that target tyrosine residues^12, 13^. These new approaches are fast and reactive under mild conditions^13^. Tyrosine is commonly found in proteins, comprising on average 3.2% of residues in native protein structures, on average^14, 15^. The phenolic ring of tyrosine is partly hydrophobic and polar. Tyrosine residues can be found at the protein surface or buried in the hydrophobic core^13, 15, 16^. Protein surfaces that bind small molecules, nucleic acids, or protein partners are frequently enriched in tyrosine^13, 17, 18^. These properties contrast with the amino acids lysine and cysteine, which are the most frequent targets for chemical modification^19, 20^. Lysine is highly abundant but largely restricted to protein surfaces, owing to its positively charged sidechain. Cysteine is rare in proteins and is typically buried because of its tendency to form disulfide bonds. Lysine and cysteine residues are constrained to occupying similar positions in alternative protein conformations, irrespective of significant global changes to the protein structure.

Recent advances in targeting tyrosine include labeling through a click-like reaction using 4-phenyl-3H-1,2,4-triazole-3,5(4H)-dione (PTAD)^21, 22^. Here, we investigated the use of PTAD to indicate changes in protein structure. We applied PTAD conjugation to folded and unfolded proteins and analyzed the relative differences in labeling with mass spectrometry. Our goal was to demonstrate whether tyrosine modification by PTAD might provide a useful tool to distinguish protein conformations. Our results may enable future adaptions of this approach to probe protein structure in cases where traditional methods are limited.

## RESULTS

### Conjugation of tyrosine and peptides by PTAD

We assessed a previously established method of conjugating tyrosine with PTAD^21, 22^. We validated the reaction for a tyrosine analogue, 3-(4-hydroxyphenyl)propionic acid (**Supplemental Spectra 1**), by comparing reactants and products through ^1^H-NMR. After a 15-minute incubation of PTAD (1.15 mM) and propionic acid (0.5 mM), we observed peaks to indicate a new species in the ^1^H-NMR spectra, which matched previously reported spectra of the conjugated product (arrows in **Figure 1A**)^23, 24^. To further distinguish the product spectra from the propionic acid reactant, we repeated the 15-minute reaction for the same sample four times, until the unreacted propionic acid signals were nearly abolished, and product signals remained. We observed that the reaction occurred quickly, even in mild aqueous conditions, as indicated in previous reports^21, 24^.

**Figure 1.**
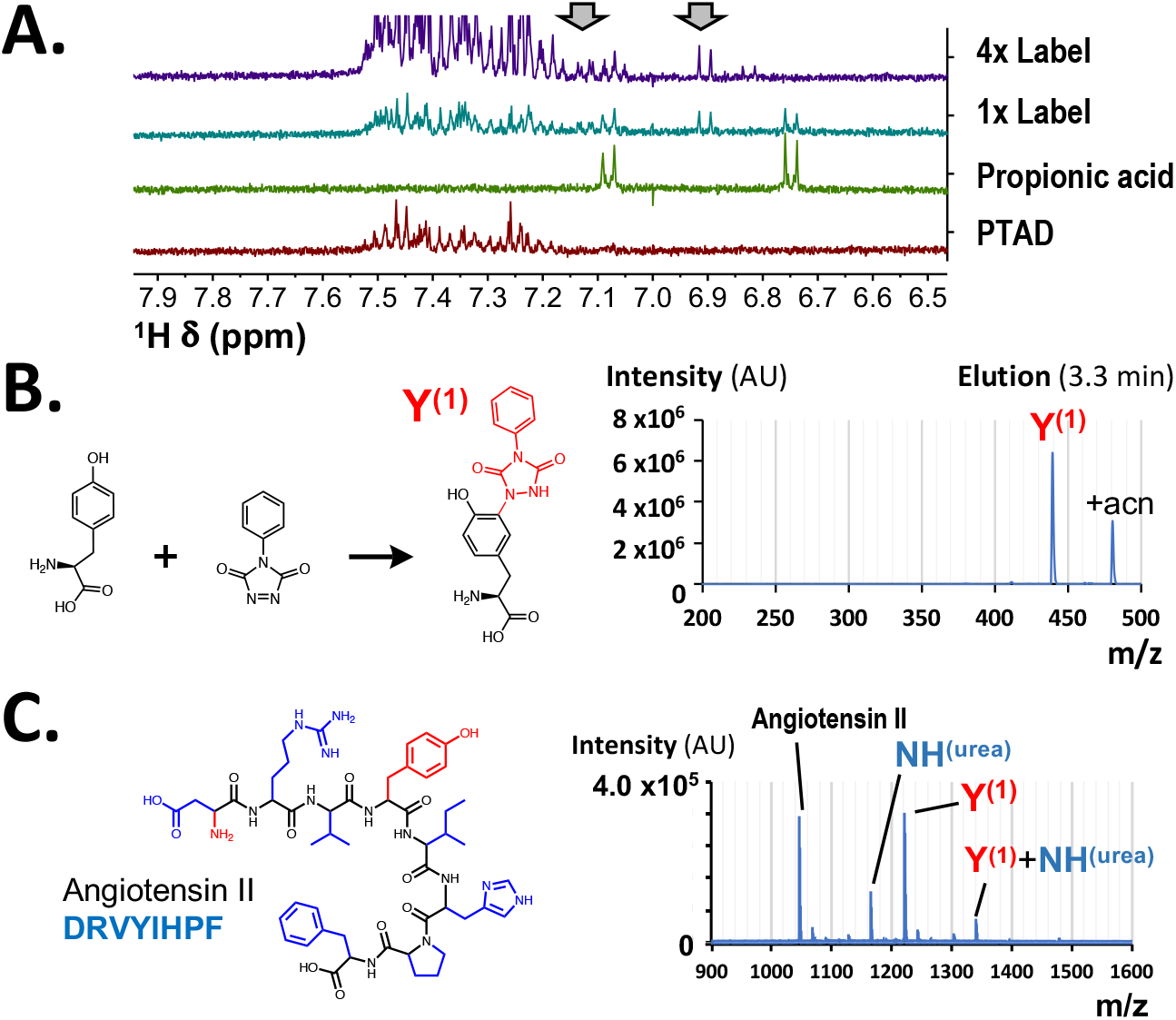
Products of PTAD reaction with tyrosine. **(A)** ^1^H NMR spectra were compared for PTAD, propionic acid, a 15-minute conjugation reaction (1x Label), and a 15-minute reaction repeated for a single sample four times sequentially (4x Label). Arrows indicate regions where the peak of the product was expected^23, 24^. **(B)** The conjugation product of 1:1 PTAD to tyrosine at a single *ortho*-position, Y^(1)^, on the phenolic ring, was detected by UPLC-MS with the expected m/z of 440 Daltons. **(C)** Products of Angiotensin II were observed with MALDI-TOF. Products shown include a PTAD conjugate with the tyrosine at position 4, Y^(1)^, and a conjugate of a phenylisocyanate to the N-terminal amine, NH^(urea)^. AU indicates arbitrary units for MS intensities. All chemical structures are shown in **Supplemental Diagram 1**.

We next tested the products of conjugation with tyrosine by ultra-performance liquid-chromatography mass spectrometry (UPLC-MS). Oxidized PTAD is commercially available in powder form and can be reacted from fresh stocks. For our UPLC-MS analysis, we used reduced PTAD-azide (red·PTAD-N_3_, m/z = 262). This form can be stored in solution and was activated to PTAD-N_3_ by 1,3-dibromo-5,5-methylhydantoin (DBH) (see Methods). A 1:1 molar ratio (2.25 mM) of PTAD-N_3_ and tyrosine produced three products after 1 hour. We observed a PTAD and tyrosine conjugate, Y^(1)^, by UPLC-MS (m/z = 440, **Figure 1B**). We also observed a double-reacted product with PTAD conjugated to each *ortho* carbon of the phenolic ring, Y^(2)^ (m/z = 700, **Supplemental Figure 1A**). Previous reports have also noted modifications of amines during PTAD reactions, and short-lived conjugation to cysteine residues^23, 25^. PTAD degradation yields an isocyanate protein that is reactive to amines^23, 26^. We observed the product of this reaction at the mass expected for the phenylisocyanate conjugated to the primary amine of tyrosine, NH^(urea)^ (m/z = 384, **Supplemental Figure 1A**). Dilution of tyrosine until the PTAD (2.25 mM) was at 10 or 50-fold molar excess yielded combined products of Y^(1)^ or Y^(2)^ and the NH^(urea)^ conjugation.

Finally, we assessed the products of PTAD (m/z = 175) and a peptide, Angiotensin II. Angiotensin II is a peptide 8 amino acids long peptide, with a sequence DRVYIHPF (m/z = 1046, **Figure 1C**). In this reaction, we incubated PTAD in aqueous buffer containing 200 mM 2-amino-2-hydroxymethyl-propane-1,3-diol (Tris) to scavenge the isocyanate through its own primary amine^21, 23^. After a 1-hour incubation at room temperature, we detected the PTAD conjugate (m/z = 1221) by matrix-assisted laser desorption/ionization and time of flight (MALDI-TOF) mass spectrometry. We also observed minor products with the N-terminal amine labeled (m/z = 1165) or both the N-terminal amine and tyrosine conjugated (m/z = 1340). To further evaluate quenching of this side reaction, we repeated Angiotensin II labeling in 10 mM, 100 mM, or 1 M Tris and observed reduced or abolished N-terminally labeled products with increasing Tris concentrations (**Supplemental Figure 1B**).

### BSA protein remains folded during PTAD conjugation

We hypothesized that the reactivity of tyrosine would depend on its position in a folded protein. However, protein unfolding due to reaction conditions might limit the ability to distinguish alternative protein conformations on the basis of their patterns and levels of PTAD labeling. To explore the stability of a protein during labeling with PTAD, we chose to investigate the well-folded protein bovine serum albumin (BSA, MW = 66 kDa), comprising of 607 amino acids and 21 tyrosine residues.

Two approaches providing a straightforward assessment of protein folding are size exclusion chromatography (SEC) and circular dichroism (CD) spectroscopy. We compared the profiles for PTAD-labeled BSA with those for BSA unfolded by titrating amounts of urea^27^. Circular dichroism (CD) spectroscopy revealed a midpoint concentration for BSA unfolding as 4.6 ± 0.1 M urea (**Supplemental Figure 2A**). Next, we compared the CD spectra for BSA with and without PTAD labeling. Interference by PTAD absorbance required the spectra to be collected at lower protein concentrations. Nevertheless, the CD spectra of PTAD-labeled BSA did not indicate protein unfolding (**Supplemental Figure 2B**).

We then used SEC, as a complement to CD spectroscopy, to compare folded or unfolded BSA and PTAD-labeled protein. During SEC, a single peak of BSA was observed at an elution volume near 14.5 mL. BSA denatured in 4, 6, or 8 M urea eluted earlier with broader peaks, thus indicating a more extended structure (**Figure 2A**). We labeled BSA with equimolar PTAD (2.25 mM) to tyrosine (121 μM BSA = 2.5 mM tyrosine). SEC revealed a single peak corresponding to BSA at 14.5 mL and a second peak near the end of the column volume due to the absorbance of the unreacted PTAD, which is reduced in water^21, 25^. The SEC profile for BSA eluting near 14.5 mL was not changed by PTAD conjugation (**Figure 2B**, solid red line). A second peak seen near 21 mL for the labeled BSA samples was confirmed by SDS-PAGE to be not protein but absorbance at 280 nm of unreacted PTAD (“Free label”) (**Supplemental Figure 2C**). PTAD and DBH stock solutions are often in organic solvents that may influence protein structure. We did not observe protein unfolding for labeling while the percentage of acetonitrile/H2O was at or below 5%. Reactions with 20% acetonitrile/H2O did produce a broad peak consistent with unfolded protein (**Figure 2B**, dashed red line). We also tested an expedient method of detecting PTAD labeling through a conjugated fluorophore. BSA labeled with PTAD-N_3_ click conjugated by DBCO-Cy5 did not appear to be unfolded to analysis in analysis by SEC (**Supplemental Figure 2D**).

**Figure 2.**
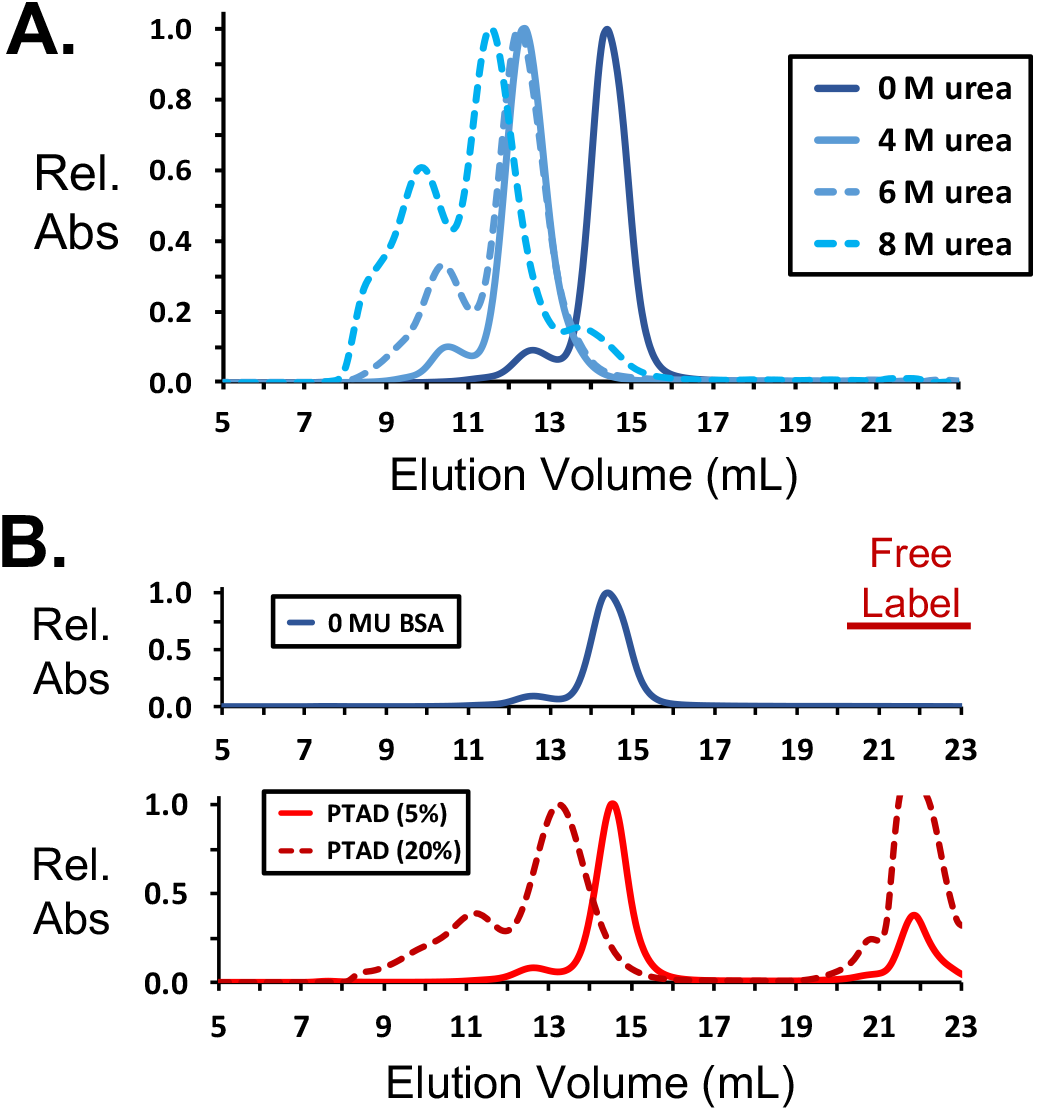
PTAD conjugation does not abolish protein structure. (**A**) SEC of BSA was monitored by UV absorbance and with titrating amounts of urea. The BSA elution was shifted after unfolding in urea. “Rel. Abs” in all SEC plots represents the relative absorbance measured at 280 nm. (**B**) PTAD conjugated BSA elution (solid red line) completely superimposed over that of folded BSA (blue line). Labeling of BSA in 20% ACN unfolded the protein, thus causing a shift in its elution (dashed red line). The peaks near 22 mL are indicated as free label, which results from strong absorbance at 280 nm for unreacted PTAD. An SDS-PAGE gel of selected fractions for the PTAD (5%) sample is found in **Supplemental Figure 2C**.

### Alternative folded states for BSA are distinctly labeled by PTAD

We next used liquid chromatography with tandem mass spectrometry (LC-MS/MS) to map PTAD labeling to individual tyrosine residues in BSA. We were able to detect PTAD labeling at all previously published sites in BSA except Y393, for which as few as 2 peptides were observed in some replicates^22^. We also found PTAD labeling at several residues not previously reported: Y173, Y179, Y180, and Y520^21, 22, 28, 29^. We interpreted this finding as the result of recent improvements in sensitivity in LC-MS/MS technology. We quantified the ratio of labeled to unlabeled tyrosine (L/U) for 13 tyrosine residues (N = 4, **Supplemental Figure 3A-B**). Only single PTAD conjugations to tyrosine were observed for residues positioned at or near the protein surface (**Figure 3A**). PTAD was quantified for 11 tyrosine residues. Phosphorylation was detected for three tyrosine residues (**Supplemental Figure 3A**). Y286 had the highest phosphorylation signals and was not found to be labeled by PTAD. Y520 occupied a disordered region of the protein with an adjacent phosphorylated threonine, T519, and was also not found to be labeled by PTAD (**Supplemental Figure 3A**).

**Figure 3.**
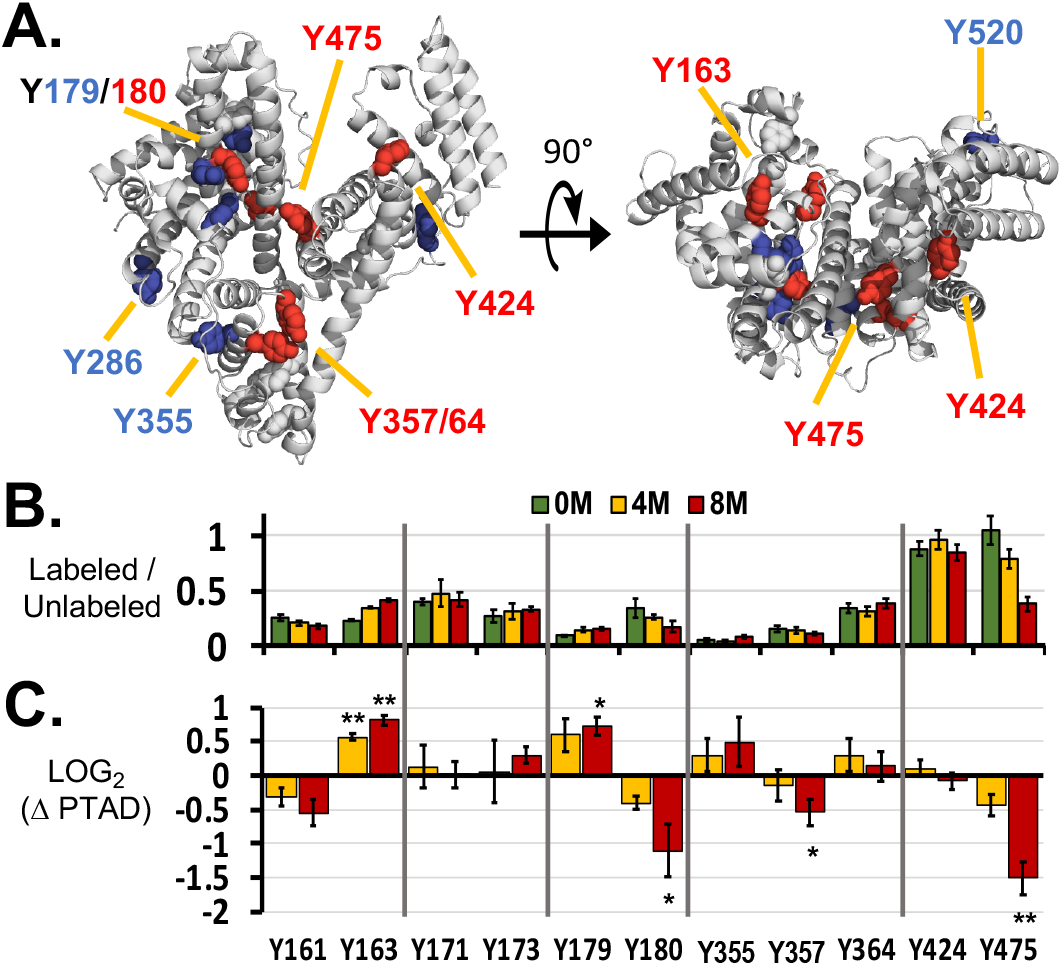
Quantitative analysis of PTAD labeling for BSA. (**A**) Two views of BSA (PDB: 3V03) are shown with tyrosine residues represented as spheres. Tyrosine residues are colored according to PTAD labeling was observed more (red) or less (blue) than expected. (**B**) The ratio of labeled to unlabeled (L/U) tyrosine is shown for each residue in native BSA, 0 M urea, and BSA incubated overnight in 4 M or 8 M urea. (**C**) The LOG_2_ of the fold change in L/U is also shown for BSA incubated in 4 M and 8 M urea, as compared with the native structure in 0 M urea. Error bars are standard error of the mean (N = 4 for all treatments). * p < 0.05, and ** is p < 0.005, Student’s t-test assuming equal variances.

We used Angiotensin II to confirm that the presence of urea did not interfere with labeling by PTAD. The ratio of labeled to unlabeled peptide observed by MALDI-TOF was not changed for peptides labeled with or without an overnight incubation in 4 M or 8 M urea (**Supplemental Figure 3C**). In the same way, BSA was incubated overnight with 4 M or 8 M urea and labeled (N = 4 for each treatment). LC-MS/MS analysis for these samples was performed in parallel with analysis of the native (0 M urea) BSA samples, and levels of PTAD labeling were compared (**Figure 3B**). We compared the L/U ratio for each tyrosine position relative to its average in the folded (0 M urea) protein. The LOG_2_ of the fold changes are plotted to show positive and negative changes on a similar scale (**Figure 3C**). Levels of PTAD were determined to be unchanged for residues Y171, Y173, Y364, and Y424 between the alternative conformations. The largest increase in PTAD labeling was found for Y163 (p < 0.001 for 8 M urea; Student’s t-test). The tyrosine residues with the lowest L/U ratios, Y179 and 355, were also labeled more frequently in 8 M urea than in the native state (p = 0.019 and p > 0.05, respectively, for 8 M urea; Student’s t-test).

For the 8 M urea samples, the residue with the greatest decrease in PTAD levels was Y475, which was the most labeled residue in the native (0 M urea) state (**Figure 3C**). A significant decrease in PTAD conjugation was also found for Y180 and Y357 (p = 0.009 and 0.0034, respectively, for 8 M urea; Student’s t-test). The decrease in levels of PTAD was greater for 8 M than 4 M urea samples. In contrast to tyrosine, highly charged lysine residues are predicted to be similarly solvent exposed in both the folded and unfolded states, and therefore no change in lysine modification by phenylisocyanate was expected. Phenylisocyanate (m/z = 121) conjugation was observed for 16 lysine residues – twice as many as was previously reported^21, 23^. However, this modification showed little change between the alternative conformations produced in 4 M or 8 M urea (**Supplemental Figure 3C**).

### PTAD conjugation is sensitive to protein structure

We compared the locations of tyrosine residues in the solved BSA structure (PDB: 3V03) with the levels of PTAD found conjugated at that position. We expected to find a simple relationship between the level of PTAD labeling and solvent exposure of a given residue. We used the molecular visualization tool PyMOL to calculate the solvent accessible surface area for each tyrosine in BSA. A simple relationship appeared to exist for some residues. The L/U ratios for residues Y355, Y357, and Y364 (3/113, 15/101, and 117/407, respectively) were linearly correlated with the percentage of surface area exposed to solvent (2, 13, and 21%, **Figure 4A**). The residues calculated to be the most solvent exposed, Y424 and Y475, were found to have the highest L/U ratios (**Figure 4B**).

**Figure 4.**
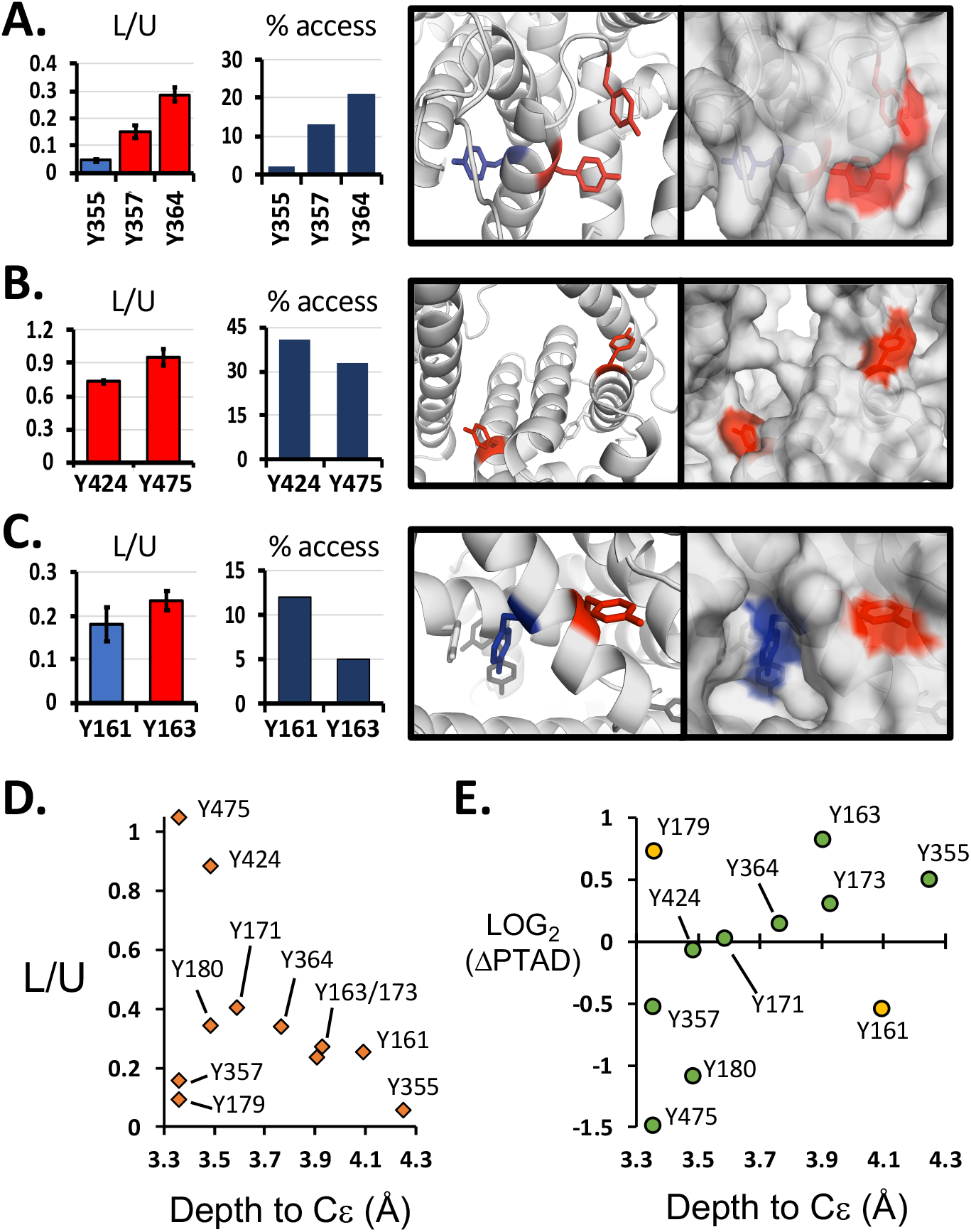
Effects of local structure on PTAD labeling. The ratio of labeled to unlabeled tyrosine residues (L/U) was measured. Residues enriched in PTAD are shown in red in the left bar graph and structures. Those with relatively low ratios are shown in blue. In addition, the percentage accessible surface area calculated for a solved structure for BSA (PDB: 3V03) is displayed. **(A)** For Y355, Y357, and Y364, the amount of PTAD labeled tyrosine increased as the percentage accessibility increased. **(B)** Y424 and Y475 were found to have the highest L/U ratios and percentage accessibility. **(C)** Y163 was predicted to have considerably lower solvent accessibility but its enrichment in PTAD labeling was approximately equal to that of Y161. In the structure, Y163 can be seen to have its *ortho* position oriented toward the surface.

The extent of PTAD labeling could not be strictly inferred from solvent exposure in the solved BSA structure. Residues Y161 and Y163 were measured to have similar levels of PTAD labeling (49/326 and 62/326) but markedly differed in their surface area predicted to be solvent exposed (**Figure 4C**). Y161, was more exposed (12%) in the crystal structure than the nearby residue, Y163 (5%). However, we also noted that the phenolic ring of Y163 was oriented with the *ortho* positions facing toward the solvent. The exposed surface of Y161 was from the backbone and *meta* positions, and the hydroxyl and *ortho* positions were mostly buried (**Figure 4C**, right). On the basis of this finding, we hypothesized that solvent exposure for a tyrosine residue may not completely predict the accessibility to PTAD conjugation^12^.

A complex relationship between labeling and solvent exposure was observed for Y171, whose L/U ratio was greater than that of the nearby residue Y173 (44/143 and 32/143), although both residues had similar solvent exposure (**Supplemental Figure 4A**). The residues Y179 and Y180 diverged considerably in their L/U ratios relative to their solvent exposure (**Supplemental Figure 4B**). We found that the phenolic ring for the more exposed residue, Y179, lies nearly parallel to the protein surface, whereas that of Y180 is perpendicular, with the *ortho* position most exposed. Another potential influence on Y180 labeling was the nearby basic residue, K187, which may alter local pH (**Supplemental Figure 4B**).

Using another approach to relate BSA structure to tyrosine modification, we investigated the depth of carbon atoms Cε1 and Cε2, at the *ortho* positions of the tyrosine sidechain. We calculated each atom’s depth from the solvent accessible surface in PDB 3V03 by using the Euclidean Distance Transform based Surface program, EDTSurf^30, 31^. We averaged the depth measured for Cε1 and Cε2 and compared the values to the levels of PTAD labeling. In the native BSA structure, we found that most tyrosine L/U ratios related linearly to atom depth, and the most labeling was found for residues nearest to the native protein surface (**Figure 4D**). Exceptions were Y179 and Y357, which had less PTAD label than for other residues similarly positioned near the protein surface.

We also compared the distance from the native protein surface and the differences between the native structure and the conformation in 8 M urea. Most tyrosine residues near the surface were labeled less in 8 M urea relative to the native fold (**Figure 4E**). Residues at greater distances from the surface were found to have higher L/U ratios in the conformation unfolded in 8 M urea. Exceptions were found for Y179 and Y161. For Y179, the low L/U ratio observed in the native protein (**Figure 4D**) was similar to that of its neighboring residue Y180 for the 8 M urea conformation. The solved crystal structure of BSA did not suggest a simple explanation for the decrease in Y161 L/U ratios in the 8 M urea conformation. We interpreted the simplest explanation to be that at position Y161, the solved structure may deviate from the predominant residue position in solution. From the 11 tyrosine residues quantified with LC-MS/MS, decreased labeling was found in the unfolded protein for most residues whose average Cε depth was less than 3.5 Å. Most residues whose average Cε depth exceeded 3.8 Å were found to show increased labeling in the unfolded protein.

We found that PTAD effectively labeled tyrosine residues in folded or unfolded protein conformations. The protein maintained much of its shape after labeling with PTAD, according to SEC and CD spectroscopy analysis. Importantly, we demonstrated substantial changes in tyrosine labeling by PTAD that could be used to distinguish alternative protein conformations for native or unfolded BSA. These results demonstrate that tyrosine conjugation can provides a chemical fingerprint that can indicate differences in protein conformation.

## DISCUSSION

In agreement with findings from earlier reports, we found that PTAD efficiently conjugates tyrosine residues in short time periods under standard conditions to maintain protein structure or activity^21, 22, 24^. We could distinguish three previously reported products of PTAD conjugation: a single phenolic addition to tyrosine, a double addition, and an isocyanate reaction with a primary amine (**Figure 1**). Labeling of the protein BSA was mapped and quantified with LC-MS/MS for 11 tyrosine residues distributed throughout the protein (**Figure 3B**). The pattern and amount of labeled tyrosine were distinct for each conformational state in 0 M, 4 M, or 8 M urea. In most cases, a comparison between the amount of labeled tyrosine and its position in the solved crystal structure for BSA suggested a simple relationship based on relative solvent accessibility. This finding suggests that tyrosine conjugation by PTAD could be developed as a tool to aid in investigations of protein structure-function relationships.

Structural studies and previous investigations of tyrosine modifications in BSA suggest that tyrosine residues vary in their accessibility to solvent^9, 10, 32^. These studies suggest that analysis of PTAD labeling would also find the presence of PTAD varies according to solvent accessibility of each tyrosine. The extent of residue exposure to solvent is among the simplest measurements that can be extracted from a solved structure by using molecular visualization software, such as PyMOL. We found that labeling and solvent exposure often appeared to correlate, on the basis of LC-MS/MS analysis (**Supplemental Figure 3A**). However, some tyrosine residues did not follow a simple trend.

A possible explanation for the unexpected levels of tyrosine labeling may be protein movement in solution^33^. Tyrosine residues can undergo ring flipping despite being well packed in a folded protein^34, 35^. The crystal structure of BSA suggested substantial variation in the orientation of tyrosine sidechains relative to the protein surface (**Figure 4, Supplemental Figure 4**). We observed that the depth from the protein surface, averaged for carbons Cε1 and Cε2 at the *ortho* positions of tyrosine, offered a better explanation for the relative levels of tyrosine conjugation by PTAD (**Figures 4D–E**). Tyrosine residues with their *ortho* carbons close to the surface were preferentially reacted with PTAD, but this enrichment was lost with protein unfolding. Similarly, tyrosine residues whose *ortho* carbons were buried were occluded in the folded protein but more easily labeled in the unfolded state (**Figure 4E**).

By LC-MS/MS, we observed 13 tyrosine residues in BSA, of which 11 were modified by PTAD. A caveat was that some residues not modified by PTAD were already modified through phosphorylation or were near phosphorylated residues. PTAD is not expected to react with phospho-tyrosine^13^. This aspect may expand the ability of this approach to distinguish protein states. Nine of the labeled tyrosine residues (82%) followed a trend in which the amount of tyrosine label was reduced as the distance from the protein surface increased. Notably, this approach was sensitive, even in distinguishing between partially and fully disordered states (**Figure 3C**). Therefore, this may potentially be used for this approach to analyze intrinsically disordered proteins. Future studies across a wider range of proteins may reveal whether this conjugation-based approach can reveal novel aspects of protein structure that cannot be predicted or inferred in analyses with standard approaches, such as NMR, X-ray crystallography, and hydrogen-deuterium exchange^3, 19, 36, 37^.

In conclusion, we consider quantitative analysis of protein structure by tyrosine conjugation to be a practical method for protein analysis. In this study, alternative protein conformations for BSA were easier to distinguish according to tyrosine labels than lysine labels. Through the use of new reagents, such as PTAD, opportunities may arise to apply a conjugation-based approach to probing protein structure under circumstances preventing analysis through standard structural biology approaches. In the future, the application of this approach to whole cell proteomics and investigations of tyrosine-rich low complexity proteins may shed light on the extent to which this technique can reveal new protein biology.

## METHODS

### Materials

PTAD conjugation reagents were purchased commercially from Sigma-Aldrich (Germany) and used without further purification: 4-phenyl-3H-1,2,4-triazole-3,5(4H)-dione (PTAD, cat. #42579), 4-(4-(2-azidoethoxy)phenyl)-1,2,4-triazolidine-3,5-dione, N3-Ph-Ur for e-Y-CLICK (PTAD-N_3_, cat. #T511552), 1,3-dibromo-5,5-dimethylhydantoin (DBH, cat. #157902), DBCO-Cy3 (cat. #77366), DBCO-Cy5 (cat. #777374), and tyrosine (cat. #T3754). Tris-HCl was purchased from Goldbio (cat. #T-400-5). Urea was purchased from Invitrogen (Carlsbad, CA). Angiotensin II was purchased commercially from Sigma-Aldrich (cat. #A9525). Bovine serum albumin (BSA) was purchased from VWR (cat. # 97062-508). Chemical structures are provided in **Supplemental Diagram 1**.

### Nuclear Magnetic Resonance Spectroscopy

For analyses using ^1^H NMR, 3-(4-hydroxyphenyl)propionic acid (20 mM) was dissolved in MilliQ filtered water (20 mM) and PTAD (115 mM) was dissolved in acetonitrile. Stocks were combined with 0.5 mM propionic acid and 1.15 mM PTAD in 1 mL of water and 4% acetonitrile. Reactions were incubated for 1 hour at room temperature. The 4x labeled samples were produced by repeating the addition of PTAD into the reaction four times at 0, 15, 30, and 45 minutes. The final concentration of PTAD after the fourth addition was 0.5 mM propionic acid and 4.6 mM PTAD in water and 4% acetonitrile. Propionic acid, PTAD, and products from the two reaction conditions were lyophilized overnight and resuspended in D_2_O. ^1^H NMR spectra were collected with a Bruker AVIII-400. Full spectra are shown in **Supplemental Spectra 1**. Spectra were analyzed by comparison to published spectra, thus confirming that the products were consistent those previously reported^23, 24^.

### UPLC-MS Analysis

Tyrosine stocks were dissolved in 1 M HCl in MilliQ filtered water. Immediately before use, red·PTAD-N_3_ (100 mM in DMF) and DBH (98 mM in DMF) were mixed (50:50, v/v) until a color change to cranberry red was observed (< 1 minute), indicating the formation of active PTAD-N_3_. PTAD-N_3_ and tyrosine were combined for final concentrations of 2.25 mM each in acetonitrile and water (50:50, v/v). The combined solution was vortexed and incubated for 1 hour at room temperature. For reactions with a 10- and 50-fold molar excess of PTAD-N_3_ to tyrosine, the PTAD-N_3_ concentration was held constant at 2.25 mM, and the tyrosine concentrations were 0.225 mM or 0.045 mM. Reaction products were analyzed with UPLC-MS by injection of 20 μL of the sample into an LCMS-2020 (Shimadzu) with a C18 column (Onyx monolithic C18, 50 x 2.0 mm). Spectra were collected over the course of a 4-minute elution column elution with a linear acetonitrile gradient of 5–20% during the first 3 minutes.

### MALDI-TOF Analysis

Stock solutions of 5 mg/mL Angiotensin II were made from powder dissolved with mass spectrometry grade water. Stocks of 45.6 mM PTAD were dissolved in acetonitrile. Reactions comprising 0.5 mM Angiotensin II, 10 mM PTAD, and 200 mM Tris-HCl (pH 7.4) in mass spectrometry grade water were incubated for 1 hour at room temperature. Reactions were also performed with 10 mM, 100 mM, or 1 M Tris-HCl (pH 7.4). To test the effects of urea, we incubated 5 mg/mL Angiotensin II with 0, 4, or 8 M urea overnight before labeling as described above in 100 mM Tris-HCl (pH 7.4).

Peptides were analyzed by MALDI-TOF with a Bruker autoflex speed LRF (Bruker Daltonics, Germany). The matrix solution used was composed of α-cyano-4-hydroxycinnamic acid (Bruker Daltonics catalog #28166-41-8) in H2O/ACN (50:50, v/v) containing 0.1% TFA. Samples were combined with matrix solution (1:10, v/v), of which 1μL was spotted onto an MTP 384 target plate steel TF (Bruker, Part. #74115). MALDI-TOF was internally calibrated with Peptide Calibration Standard II (Bruker Daltonics, Cat. #8222570). Each sample received up to 2000 laser pulses.

### CD Spectroscopy

To monitor changes in BSA structure, we incubated 1 mg of BSA overnight in 1 mL of 0–10 M urea in water. CD analysis was performed with an OLIS DSM-20 CD spectrometer and a 1 mm pathlength cylindrical cell (Hellma, cat. #121.000-QS). CD spectra were averaged from three replicates. For labeling, 500 μL reactions were prepared with 3 μM BSA (60 μM tyrosine), 3 mM PTAD, and 500 mM Tris-HCl (pH 7.4) and incubated for 1 hour at room temperature. Afterward, unreacted PTAD was removed by dialysis overnight in 1 L of 500 mM Tris (pH 7.4) with Slide-A-Lyzer™ G_2_ dialysis cassettes (10 K MWCO, Fisher Scientific, #87729). CD spectra were averaged from two or three replicates.

### Size-Exclusion Chromatography

BSA at a concentration of 40 mg/mL was incubated overnight in 0–8 M urea and water. Immediately before SEC injection, BSA was diluted (1:5, v/v) to a final solution in 150 mM NaCl and 40 mM Tris (pH 7.4). BSA was passed through a 0.22 mm centrifugal filter, and 500 μL (4 μg BSA) was injected into an AKTA™ PureL system (GE Healthcare Life Sciences, Chicago IL), and run over a Superdex 200 GL (10/300) column (GE Healthcare Life Sciences) in buffer consisting of 150 mM NaCl, 40 mM Tris-HCl (pH 7.4). To fluorescently label BSA, was mixed red·PTAD-N_3_ (100 mM in DMF) and DBH (98 mM in DMF) at 50:50 (v/v) until a color change to cranberry red was observed, typically after several seconds. Immediately afterward, PTAD-N_3_ (2.25 mM) was combined with 0.12 mM BSA (2.25 mM tyrosine) and 22.8 mM DBCO-Cy5. Reactions were incubated for 1 hour at room temperature. After SDS-PAGE separation, fluorescence in gels was imaged with a Chemidoc MP system (Bio-Rad) followed by staining with Instant Blue (Sigma-Aldrich, cat. #ISB1L).

### LC-MS/MS

A 5 mg/mL solution of BSA was incubated overnight with 0, 4, or 8 M urea. Labeling reactions combined 5 μM BSA (100 μM tyrosine) and 10 mM PTAD in a 15 μL total buffer volume of buffer (100 mM Tris-HCl pH 7.4, 137 mM NaCl, 10 mM Na_2_HPO_4_). After a 1-hour incubation at room temperature, electrophoresis was performed on a 7.5% SDS-PAGE gel. Gels were stained with Instant Blue (Sigma-Aldrich, cat. # ISB1L), and bands for BSA (5 μg) were excised from the gel. After gel slices were treated with Lys-C, peptides were extracted and purified by C18-based desalting, as previously described^38^.

HPLC ESI-MS/MS was performed in positive ion mode on a Thermo Scientific Orbitrap Fusion Lumos tribrid mass spectrometer fitted with an EASY-Spray Source (Thermo Scientific, San Jose, CA). All solvents of liquid chromatography mass spectrometry grade. Spectra were acquired with XCalibur, version 2.3 (Thermo Scientific). Tandem mass spectra were extracted from XCalibur ‘RAW’ files and charge states were assigned with the ProteoWizard 3.0.1 msConvert Script. The fragment mass spectra were assigned with Mascot (Matrix Science, London, UK; version 2.4) with the default probability cut-off score. Variable modifications used included: PTAD (175 Da) conjugation to tyrosine; oxidation of methionine; and phosphorylation of serine, threonine, and tyrosine. Cross-correlation of the Mascot results with X! Tandem was accomplished with Scaffold (version Scaffold_4.8.7; Proteome Software, Portland, OR).

### Structural Analysis

The number of modified, N_Labeled_, and unmodified, N_Unlabeled_, tyrosine residues was summed for each residue position for BSA and across all peptides observed in LC-MS/MS. The ratio L/U ratio for each of four LC-MS/MS experiments was calculated as:

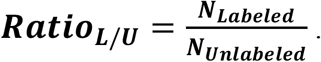

The structure for BSA, PDB:3V03, was visualized, and exposed surface areas of residues were calculated with The PyMOL Molecular Graphics System (version 2.0 Schrödinger, LLC). Accessible surface area values are calculated as a percentage of the total surface area for the residue. The depth of each atom in PDB:3V03 was calculated with the Euclidean Distance Transform based Surface program, EDTSurf^30, 31^.

## Corresponding Author

Correspondence should be addressed to Jacob C. Schwartz at.

## Author Contributions

M.M. designed and performed most experimental procedures in the manuscript, analyzed data, and interpreted results. N.K.B. performed and analyzed LC-MS/MS procedures. L.E.G. assisted designing, performing, and analyzing ^1^H NMR for PTAD reaction products. J.C.J. helped design and interpret experiments. P.R.L. designed LC-MS/MS experiments and analyzed data. J.C.S. contributed to the design of the project and experiments, data analysis, and interpretation of results. M.M. and J.C.S. collaborated in writing the first draft of the manuscript. The manuscript was written with contributions from all authors. All authors have approved to the final version of the manuscript.

## ABBREVIATIONS

BSA: bovine serum albumin
CD: circular dichroism
DBCO: dibenzocyclooctyne
DBH: 1,3-dibromo-5,5-dimethylhydantoin
LC-MS/MS: liquid chromatography and tandem mass spectrometry
NH^(urea)^: amine conjugated isocyanate
PTAD: 4-phenyl-3H-1,2,4-triazole-3,5(4H)-dione
PTAD-N_3_: 4-(4-(2-azidoethoxy)phenyl)-1,2,4-triazolidine-3,5-dione, N3-Ph-Ur for e-Y-CLICK
red·PTAD-N_3_: reduced PTAD-N_3_
SEC: size exclusion chromatography
UPLC-MS: ultra-high pressure liquid chromatography and mass spectrometry
Y^(1)^: tyrosine with single PTAD conjugation
Y^(2)^: tyrosine with two PTAD conjugations

## Table of Contents Graphic

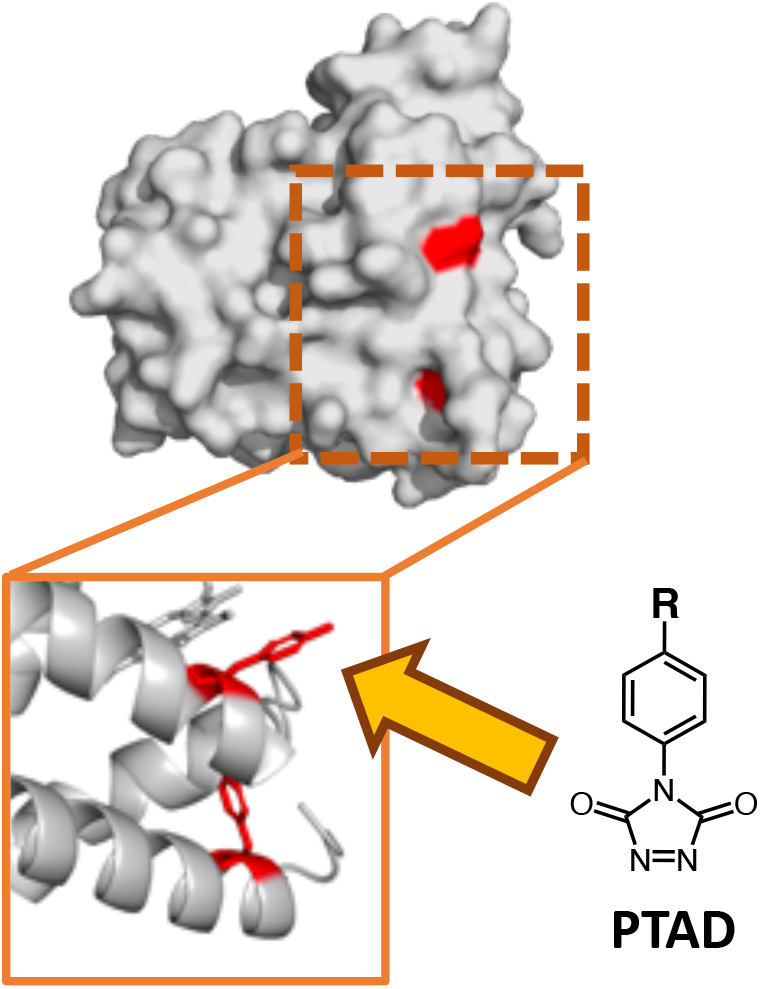

Conjugation of tyrosine in a protein is affected by local structure. Alternative protein conformations can significantly alter tyrosine accessibility. Changes in protein structure are quantifiable according to extent of tyrosine labeling observed for discrete conformational states of a protein.

## REFERENCES

1. Onuchic, J. N.; Luthey-Schulten, Z.; Wolynes, P. G., Theory of protein folding: the energy landscape perspective. Annu Rev Phys Chem 1997, 48, 545–600.

2. Anfinsen, C. B., Principles that govern the folding of protein chains. Science 1973, 181 (4096), 223–30.

3. Wang, L.; Chance, M. R., Protein Footprinting Comes of Age: Mass Spectrometry for Biophysical Structure Assessment. Mol Cell Proteomics 2017, 16 (5), 706–716.

4. Toal, S.; Schweitzer-Stenner, R., Local order in the unfolded state: conformational biases and nearest neighbor interactions. Biomolecules 2014, 4 (3), 725–73.

5. Dill, K. A.; MacCallum, J. L., The protein-folding problem, 50 years on. Science 2012, 338 (6110), 1042–6.

6. Palamini, M.; Canciani, A.; Forneris, F., Identifying and Visualizing Macromolecular Flexibility in Structural Biology. Front Mol Biosci 2016, 3, 47.

7. Uversky, V. N.; Dave, V.; Iakoucheva, L. M.; Malaney, P.; Metallo, S. J.; Pathak, R. R.; Joerger, A. C., Pathological unfoldomics of uncontrolled chaos: intrinsically disordered proteins and human diseases. Chem Rev 2014, 114 (13), 6844–79.

8. Eliezer, D., Biophysical characterization of intrinsically disordered proteins. Curr Opin Struct Biol 2009, 19 (1), 23–30.

9. Diaz-Moreno, I.; Garcia-Heredia, J. M.; Gonzalez-Arzola, K.; Diaz-Quintana, A.; De la Rosa, M. A., Recent methodological advances in the analysis of protein tyrosine nitration. Chemphyschem 2013, 14 (13), 3095–102.

10. Zhang, Y.; Yang, H.; Poschl, U., Analysis of nitrated proteins and tryptic peptides by HPLC-chip-MS/MS: site-specific quantification, nitration degree, and reactivity of tyrosine residues. Anal Bioanal Chem 2011, 399 (1), 459–71.

11. Sinz, A., Chemical cross-linking and mass spectrometry to map three-dimensional protein structures and protein-protein interactions. Mass Spectrom Rev 2006, 25 (4), 663–82.

12. Spicer, C. D.; Davis, B. G., Selective chemical protein modification. Nat Commun 2014, 5, 4740.

13. Jones, L. H.; Narayanan, A.; Hett, E. C., Understanding and applying tyrosine biochemical diversity. Molecular bioSystems 2014, 10 (5), 952–69.

14. Lee, J.; Ju, M.; Cho, O. H.; Kim, Y.; Nam, K. T., Tyrosine-Rich Peptides as a Platform for Assembly and Material Synthesis. Adv Sci (Weinh) 2019, 6 (4), 1801255.

15. Koide, S.; Sidhu, S. S., The importance of being tyrosine: lessons in molecular recognition from minimalist synthetic binding proteins. ACS Chem Biol 2009, 4 (5), 325–34.

16. Joshi, N. S.; Whitaker, L. R.; Francis, M. B., A three-component Mannich-type reaction for selective tyrosine bioconjugation. J Am Chem Soc 2004, 126 (49), 15942–3.

17. Du, H.; Hu, X.; Duan, H.; Yu, L.; Qu, F.; Huang, Q.; Zheng, W.; Xie, H.; Peng, J.; Tuo, R.; Yu, D.; Lin, Y.; Li, W.; Zheng, Y.; Fang, X.; Zou, Y.; Wang, H.; Wang, M.; Weiss, P. S.; Yang, Y.; Wang, C., Principles of Inter-Amino-Acid Recognition Revealed by Binding Energies between Homogeneous Oligopeptides. ACS Cent Sci 2019, 5 (1), 97–108.

18. Bogan, A. A.; Thorn, K. S., Anatomy of hot spots in protein interfaces. J Mol Biol 1998, 280 (1), 1–9.

19. Shannon, D. A.; Weerapana, E., Covalent protein modification: the current landscape of residue-specific electrophiles. Curr Opin Chem Biol 2015, 24, 18–26.

20. Sletten, E. M.; Bertozzi, C. R., Bioorthogonal chemistry: fishing for selectivity in a sea of functionality. Angew Chem Int Ed Engl 2009, 48 (38), 6974–98.

21. Ban, H.; Nagano, M.; Gavrilyuk, J.; Hakamata, W.; Inokuma, T.; Barbas, C. F., 3rd, Facile and stabile linkages through tyrosine: bioconjugation strategies with the tyrosine-click reaction. Bioconjug Chem 2013, 24 (4), 520–32.

22. Ban, H.; Gavrilyuk, J.; Barbas, C. F., 3rd, Tyrosine bioconjugation through aqueous ene-type reactions: a click-like reaction for tyrosine. J Am Chem Soc 2010, 132 (5), 1523–5.

23. Alvarez-Dorta, D.; Thobie-Gautier, C.; Croyal, M.; Bouzelha, M.; Mevel, M.; Deniaud, D.; Boujtita, M.; Gouin, S. G., Electrochemically Promoted Tyrosine-Click-Chemistry for Protein Labeling. J Am Chem Soc 2018, 140 (49), 17120–17126.

24. Bauer, D. M.; Ahmed, I.; Vigovskaya, A.; Fruk, L., Clickable tyrosine binding bifunctional linkers for preparation of DNA-protein conjugates. Bioconjug Chem 2013, 24 (6), 1094–101.

25. Kaiser, D.; Winne, J. M.; Ortiz-Soto, M. E.; Seibel, J.; Le, T. A.; Engels, B., Mechanistical Insights into the Bioconjugation Reaction of Triazolinediones with Tyrosine. J Org Chem 2018, 83 (17), 10248–10260.

26. Sato, S.; Nakamura, H., Protein Chemical Labeling Using Biomimetic Radical Chemistry. Molecules 2019, 24 (21).

27. Canchi, D. R.; Garcia, A. E., Cosolvent effects on protein stability. Annu Rev Phys Chem 2013, 64, 273–93.

28. Sato, S.; Hatano, K.; Tsushima, M.; Nakamura, H., 1-Methyl-4-aryl-urazole (MAUra) labels tyrosine in proximity to ruthenium photocatalysts. Chem Commun (Camb) 2018, 54 (46), 5871–5874.

29. Sato, S.; Nakamura, K.; Nakamura, H., Tyrosine-Specific Chemical Modification with in Situ Hemin-Activated Luminol Derivatives. ACS Chem Biol 2015, 10 (11), 2633–40.

30. Xu, D.; Li, H.; Zhang, Y., Fast and Accurate Calculation of Protein Depth by Euclidean Distance Transform. Res Comput Mol Biol 2013, 7821, 304–316.

31. Xu, D.; Zhang, Y., Generating triangulated macromolecular surfaces by Euclidean Distance Transform. PLoS One 2009, 4 (12), e8140.

32. Seeley, K. W.; Stevens, S. M., Jr., Investigation of local primary structure effects on peroxynitrite-mediated tyrosine nitration using targeted mass spectrometry. J Proteomics 2012, 75 (6), 1691–700.

33. Weininger, U.; Modig, K.; Akke, M., Ring flips revisited: (13)C relaxation dispersion measurements of aromatic side chain dynamics and activation barriers in basic pancreatic trypsin inhibitor. Biochemistry 2014, 53 (28), 4519–25.

34. Dreydoppel, M.; Raum, H. N.; Weininger, U., Slow ring flips in aromatic cluster of GB1 studied by aromatic (13)C relaxation dispersion methods. J Biomol NMR 2020, 74 (2-3), 183–191.

35. Rao, D. K.; Bhuyan, A. K., Complexity of aromatic ring-flip motions in proteins: Y97 ring dynamics in cytochrome c observed by cross-relaxation suppressed exchange NMR spectroscopy. J Biomol NMR 2007, 39 (3), 187–96.

36. Stuchfield, D.; France, A. P.; Migas, L. G.; Thalhammer, A.; Bremer, A.; Bellina, B.; Barran, P. E., The Use of Mass Spectrometry to Examine IDPs: Unique Insights and Caveats. Methods Enzymol 2018, 611, 459–502.

37. Bhowmick, A.; Brookes, D. H.; Yost, S. R.; Dyson, H. J.; Forman-Kay, J. D.; Gunter, D.; Head-Gordon, M.; Hura, G. L.; Pande, V. S.; Wemmer, D. E.; Wright, P. E.; Head-Gordon, T., Finding Our Way in the Dark Proteome. J Am Chem Soc 2016, 138 (31), 9730–42.

38. Parker, S. S.; Krantz, J.; Kwak, E. A.; Barker, N. K.; Deer, C. G.; Lee, N. Y.; Mouneimne, G.; Langlais, P. R., Insulin Induces Microtubule Stabilization and Regulates the Microtubule Plus-end Tracking Protein Network in Adipocytes. Mol Cell Proteomics 2019, 18 (7), 1363–1381.

