## Supplemental Data for "Discriminating changes in protein structure using PTAD conjugation to tyrosine"

**Supplemental Spectra 1:** $^{1}$H NMR spectra

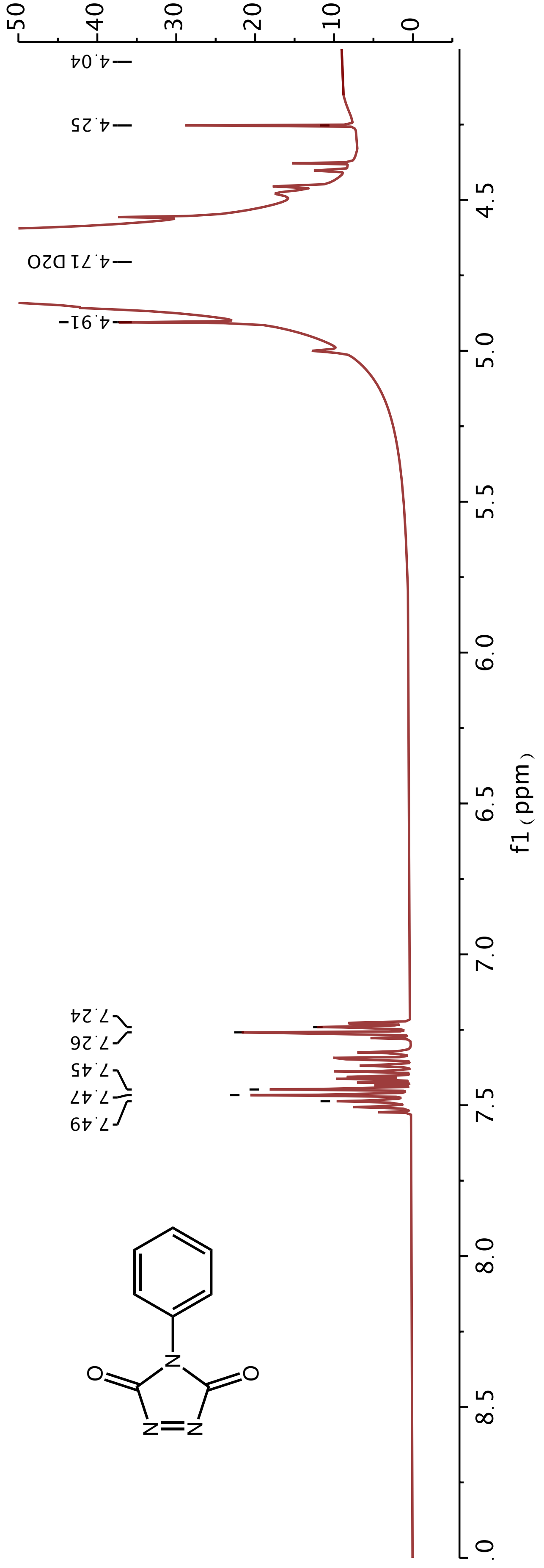


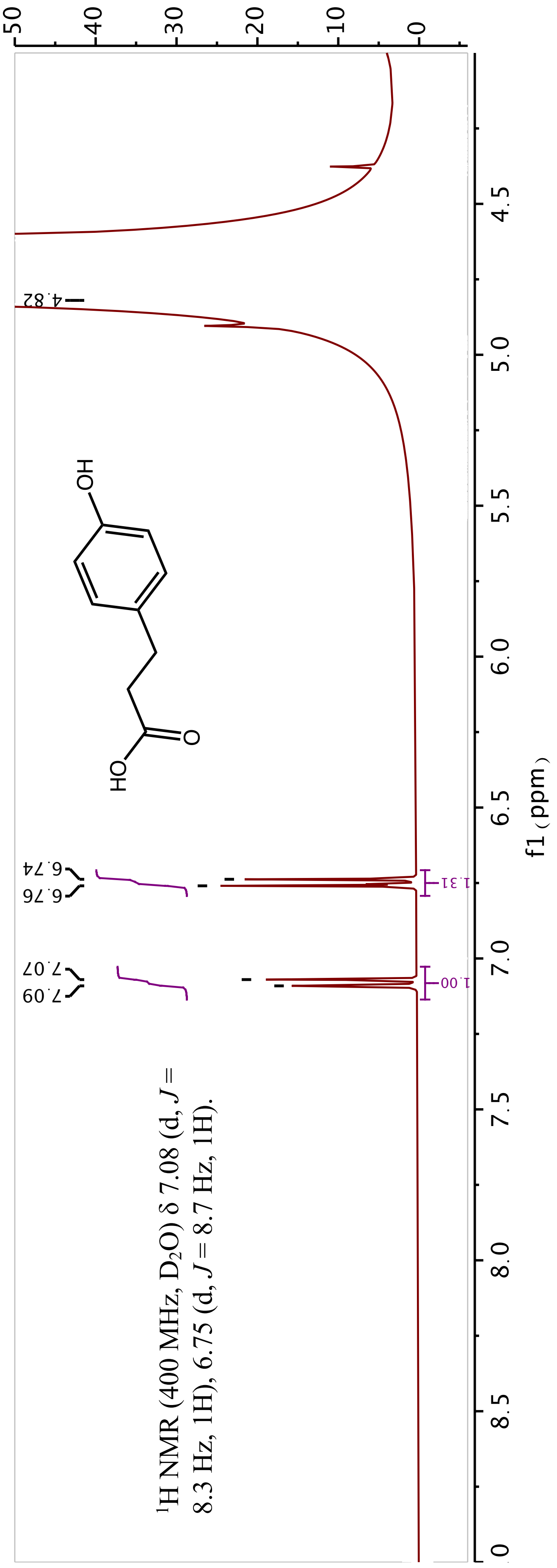

**Supplemental Spectra 1:** $^{1}$H NMR spectra

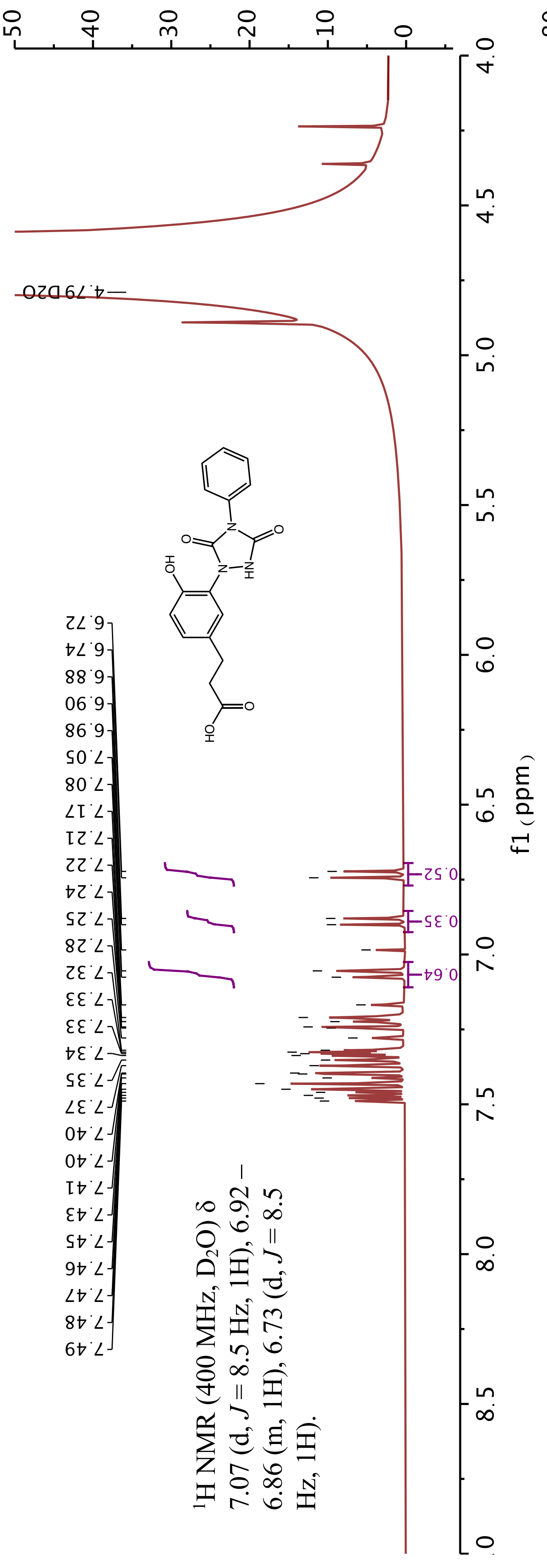


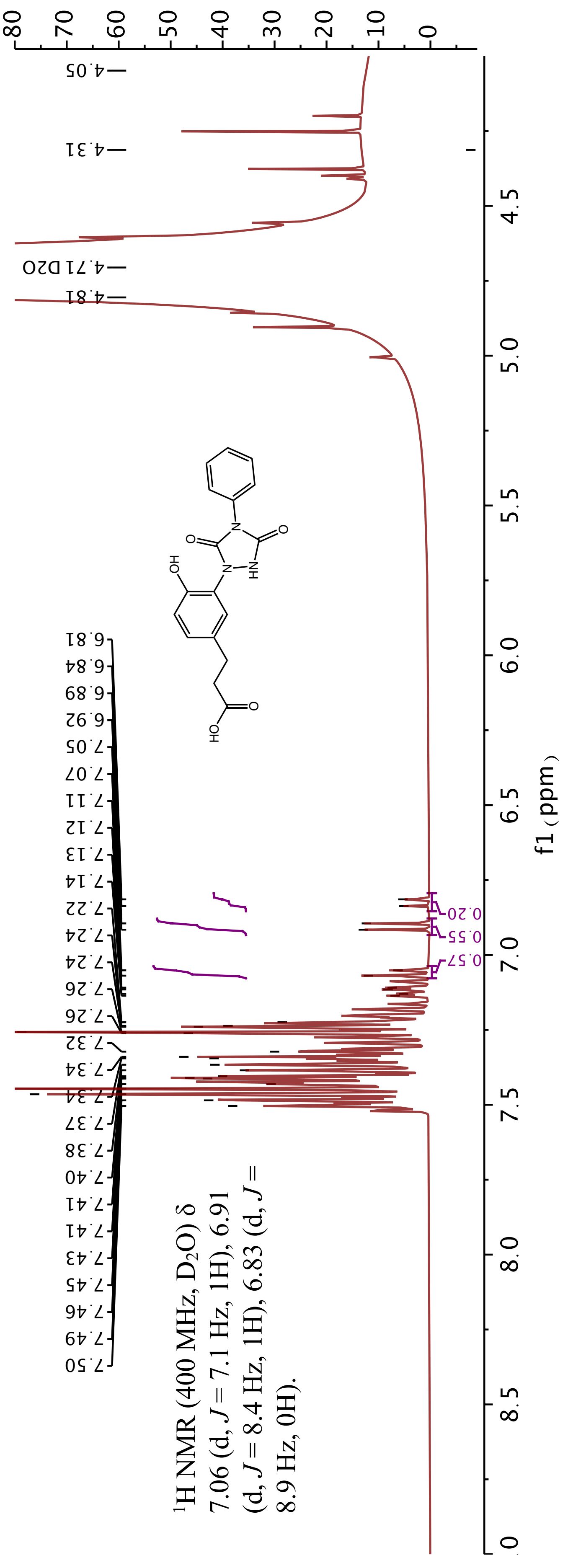

**Supplemental Diagram 1:** chemical structures for reagents

*i.*

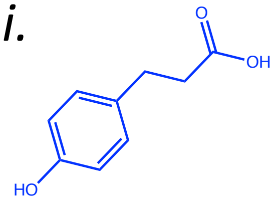


*ii.*

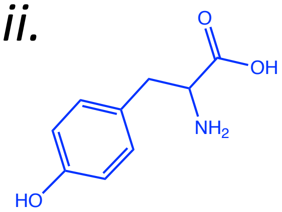


*iii.*

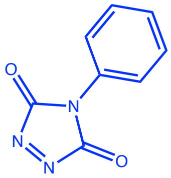


*iv.*

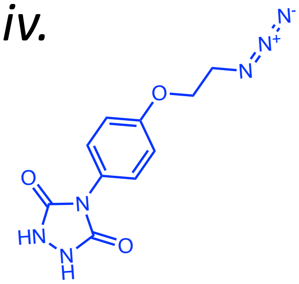


*v.*

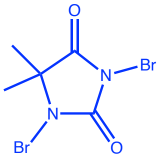


*vi.*

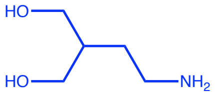


| | | |
|---|---|---|
| *i* | 3-(4-hydroxyphenyl)propionic acid | Propionic Acid |
| *ii.* | Tyrosine | Y |
| *iii.* | 4-Phenyl-3H-1,2,4-triazole-3,5(4H)-dione | PTAD |
| *iv.* | 4-(4-(2-Azidoethoxy)phenyl)-1,2,4-triazolidine-3,5-dione, N3-Ph-Ur | red•PTAD-$N_3$ |
| *v.* | 1,3-Dibromo-5,5-dimethylhydantoin | DBH |
| *vi.* | 2-amino-2-hydroxymethyl-propane-1,3-diol | Tris |

**Supplemental Figure 1:** Amine modification and Tris titration

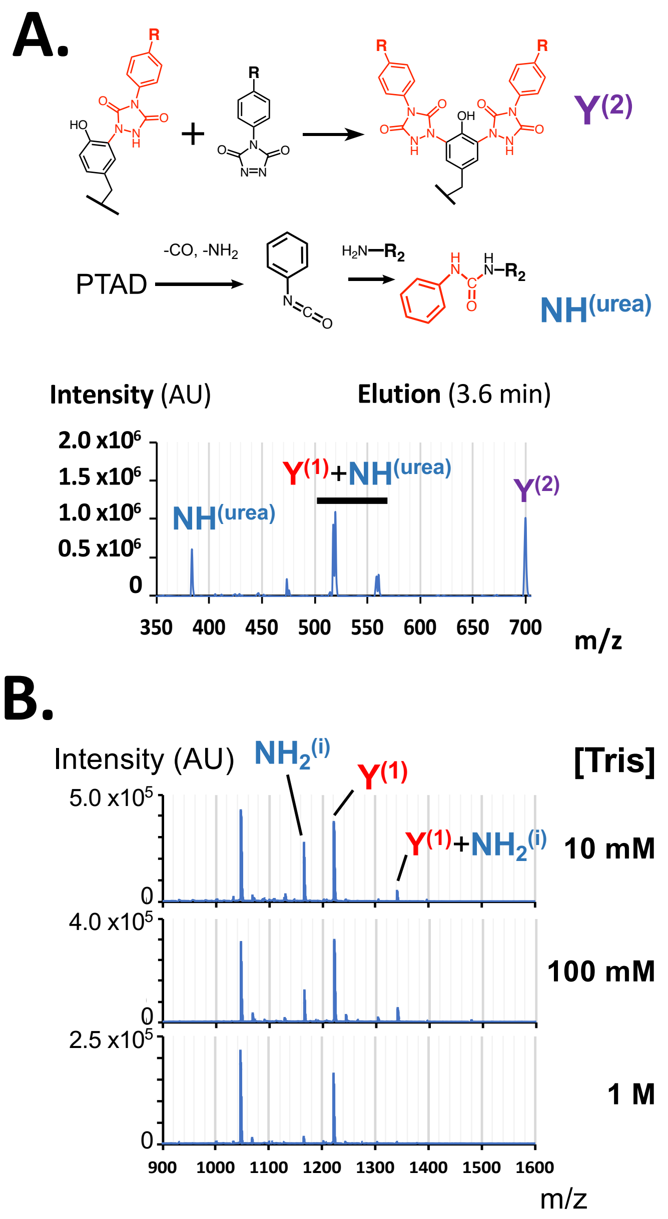


**Supplemental Figure 1**. Products of PTAD reaction with tyrosine. **(A)** Additional products are shown by UPLC-MS for PTAD and tyrosine: PTAD conjugated at both *ortho*-positions of the phenolic ring, $Y^{(2)}$, and an isocyanate degradation product of PTAD conjugated to the primary amine, $NH^{(urea)}$. **(B)** Titrating amount of Tris from 10 mM to 1 M reduces or eliminates evidence of isocyanate reaction with amines, $NH^{(urea)}$, seen by MALDI-TOF. AU indicates arbitrary units for MS intensities.

### **Supplemental Figure 2:** CD Spectroscopy and SEC data

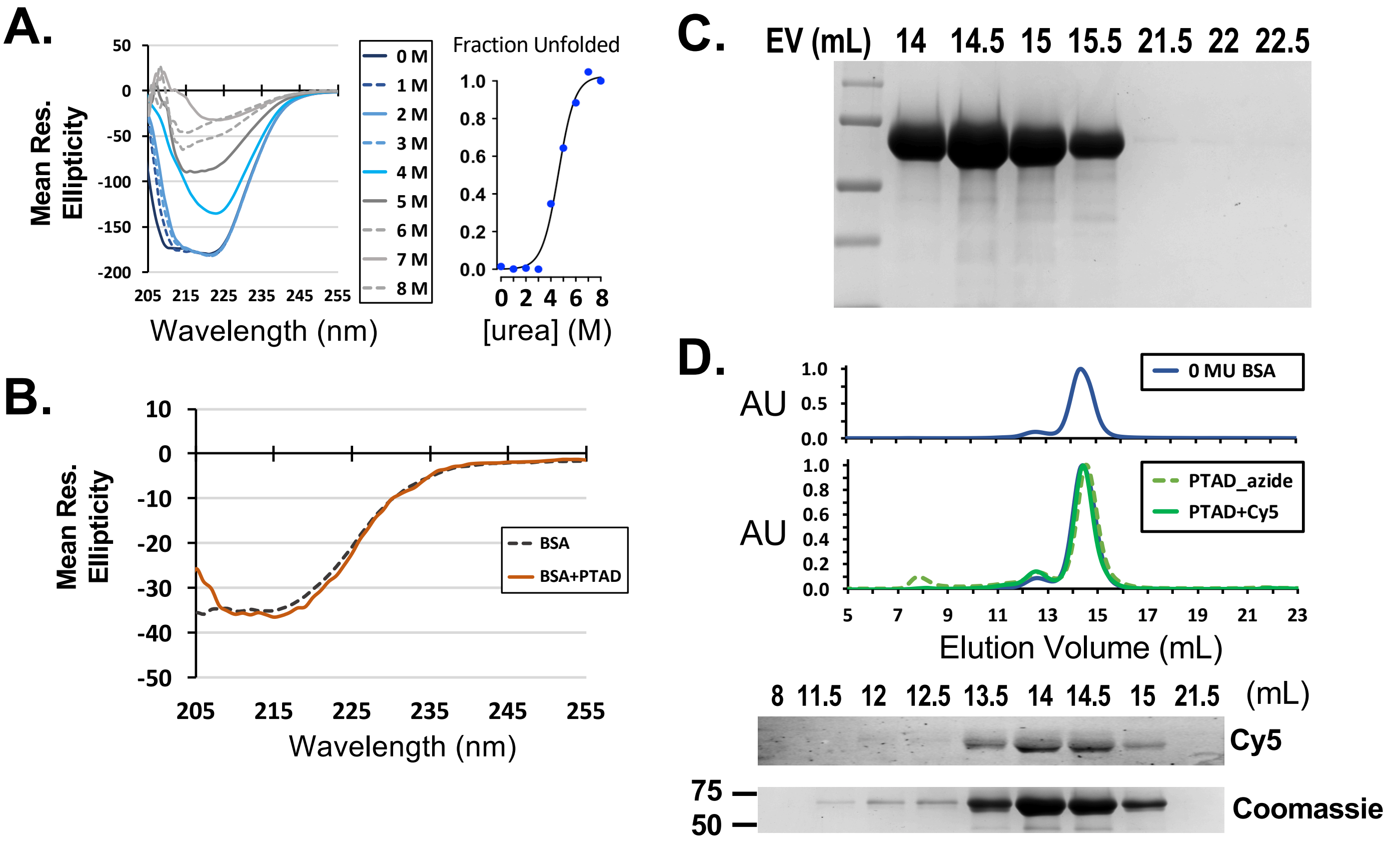


**Supplemental Figure 2**. PTAD conjugation does not abolish protein structure. (**A**) The midpoint concentration for urea to unfold BSA was determined to be 4.6 ± 0.1 M urea according to CD spectroscopy. (**B**) CD spectroscopy comparing labeled and unlabeled BSA did not indicate unfolding of the protein. Spectra were taken with a lower BSA concentration (3 μM) to mitigate interference by PTAD absorbance. (**C**) SDS-PAGE was run and stained by Coomassie for the PTAD (5% ACN) SEC experiment shown in **Figure 2B**. The samples run correspond indicated by their elution volution (EV). (**D**) Conjugations of BSA and PTAD-$N_3$ or PTAD-$N_3$ clicked with DBCO-Cy5 were indistinguishable from native BSA in SEC. Fluorescence imaging confirmed Cy5 conjugation by SDS-PAGE analysis of eluted SEC fractions. The presence of eluted BSA was also indicated by Coomassie stain.

**Supplemental Figure 3:** LC-MS/MS of BSA

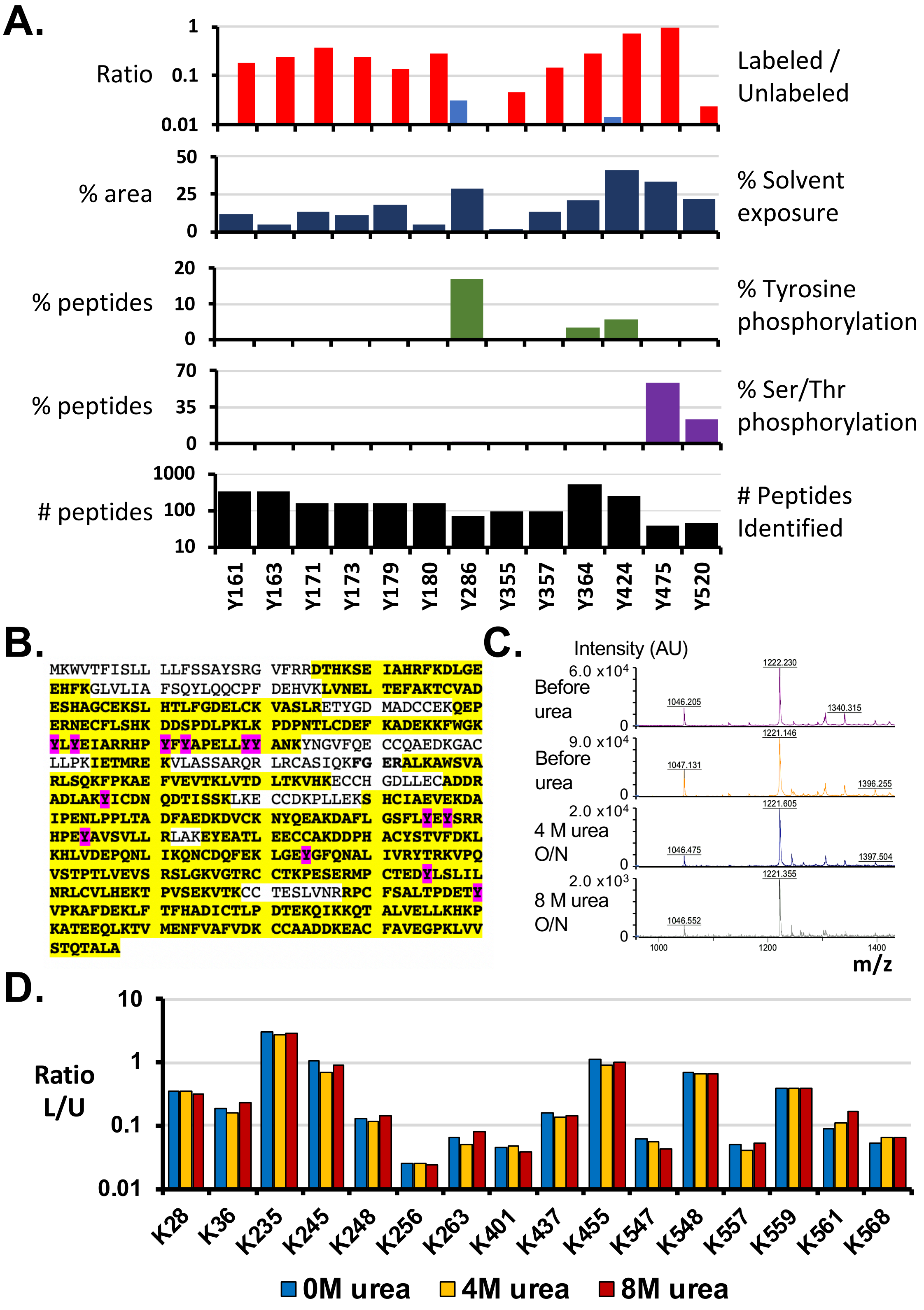

**Supplemental Figure 3**. Quantitative analysis of PTAD labeling for BSA. **(A)** The ratio of labeled to unlabeled residues is shown for BSA in 0M urea (red). Also shown are residues (light blue) called as false positives, meaning PTAD-labeled in unlabeled BSA samples. The computed solvent exposure of for tyrosine residue for BSA (PDB: 3V03) are plotted as % area (dark blue). The amount of phosphorylation the same tyrosine quantified (green) or at serine and threonine (purple) residues is shown as a percentage the total peptides containing the indicated tyrosine. Also shown is the total number of peptides detected in all replicates and containing the tyrosine indicated (black). Note that the y-axes for only the plots of L/U ratios and numbers of peptides observed are shown in log scale due to the wide range of values included. **(B)** Map of peptide coverage highlighted in yellow with tyrosines detected highlighted in pink. (**C**) MALDI-TOF analysis of Angiotensin II with or without an overnight incubation in the presence of 4 or 8 M urea. **(D)** L/U for lysine residues possessing the +119 Dalton addition of an isocyanate. Levels for this modification were predominantly unchanged under titrating amounts of urea.

#### **Supplemental Figure 4:** Analysis of PTAD labeling

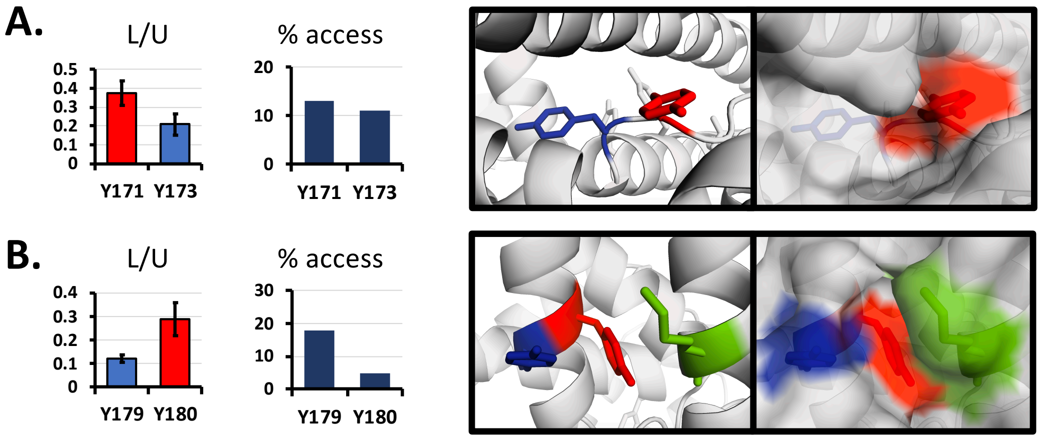


**Supplemental Figure 4**. Effects of local structure to PTAD labeling. **(A-B)** Comparisons of L/U is shown for nearby tyrosine residues L/U. The % accessible surface area for each residue is also shown (dark blue bars) for the native structure. Right, tyrosine residues are shown in sticks and their solvent exposure as a surface representation. Included in (**B**), is K187, green, which may contribute to the high L/U for Y180 despite its low % accessible surface area.
